# Multifunctional polymerization domains determine the onset of epigenetic silencing in Arabidopsis

**DOI:** 10.1101/2024.02.15.580496

**Authors:** Anna Schulten, Geng-Jen Jang, Alex Payne-Dwyer, Marc Fiedler, Mathias L. Nielsen, Mariann Bienz, Mark C. Leake, Caroline Dean

## Abstract

Cold-induced epigenetic silencing of Arabidopsis *FLOWERING LOCUS C* (*FLC*) requires the Polycomb Repressive Complex 2 and accessory proteins VIN3 and VRN5. VIN3 and VRN5 interact via head-to-tail VEL polymerization domains, but how these functionally contribute to the switch to an epigenetically silenced state remains poorly understood. Here, we determine that VIN3 VEL polymerization involves higher order nuclear VIN3 assemblies *in vivo*, promotes strong chromatin association and efficient H3K27me3 nucleation. However, we also show that the polymerization domains of VIN3 and VRN5 are not equivalent: VRN5 VEL domain is not required for silencing despite its role in physically connecting VIN3 with the PRC2 complex and VRN5 VEL is unable to functionally replace VIN3 VEL *in vivo*. Both VIN3 and VRN5 homologs are present throughout angiosperm species, suggesting a functional requirement for maintaining different polymerization modalities. This work reveals distinct roles for multifunctional polymerization domains of Polycomb accessory proteins underpinning the onset of epigenetic silencing.

## Introduction

The activity of the Polycomb Repressive Complex 2 (PRC2) is essential for the deposition of H3K27me3 as one of the hallmarks of mitotically inheritable gene silencing. Since the first identification of *cis*-regulatory Polycomb Response Elements (PREs) in Drosophila, a complex picture has emerged in which non-sequential multifactorial protein interactions with the local chromatin environment, also shaped by transcriptional activity, give rise to genome-wide Polycomb silencing patterns^1-4^. Accessory proteins that interact with the highly conserved PRC2 core complex crucially define PRC2 subcomplexes with different activities and genomic enrichment sites^5-10^. These mediate nucleation of Polycomb complexes and are required to maintain the Polycomb silenced state through cell division^10-12^. However, it is still unclear how accessory protein function facilitates the digital switch to an epigenetically stable state and ensures inheritance of the silenced state to daughter cells during DNA replication.

In the model plant *Arabidopsis thaliana*, proteins of the VEL family (VIN3, VRN5 and VEL1) have been identified as accessory proteins to a PRC2 complex with the core subunit VRN2 (one of three homologs of mammalian SUZ12) ^13,14^. Their genome-wide localization suggests a widespread role in PRC2 silencing of many loci in the Arabidopsis genome^14^. Except for VEL1, their function is essential for the epigenetic silencing of the floral repressor gene *FLOWERING LOCUS C* (*FLC)* during the winter cold, a process called vernalization^15-18^. At *FLC*, Polycomb silencing initiates via cold-induced nucleation of H3K27me3 over a small number of nucleosomes around the first exon-intron boundary of *FLC*. This confers metastable epigenetic silencing that holds the silent state for tens of cell cycles^19^. The metastable state is converted into a long-term epigenetically silenced state through the spreading of H3K27me3 across the *FLC* gene body, which occurs when the plants start to grow more rapidly following the return to ambient temperatures^20^. Nucleation, which is strongly reduced in *vin3, vrn5* and *vrn2* single mutants, can be separated from the stable spread H3K27me3 state in mutants defective in one of the PRC2 methyltransferases genes *CURLY LEAF* (*CLF*), or *LIKE HETEROCHROMATIN PROTEIN 1* required to maintain long-term silencing^21^. Studies in these mutants have shown that VEL proteins promote the stochastic PRC2-mediated nucleation of individual *FLC* alleles, whereby the fraction of nucleated *FLC* loci increases over prolonged cold of winter^22^. In other words, the slow accumulation of H3K27me3 at the 3-4 nucleosome nucleation region is the result of a low probability digital ON/OFF switch, which occurs independently at each allele^20,23^.

A key question was which features of the VEL proteins confer the ability to aid PRC2 engagement with these local sites, and to maintain epigenetic silencing through cell division? The VEL proteins share a domain architecture consisting of a tripartite plant homeodomain (PHD) superdomain, a fibronectin type III (FNIII) domain and a C-terminal VEL domain^24,25^ (Fig. 1A). Early data assumed they may be functionally similar, with one protein being cold-induced (VIN3) and the others (VRN5, VEL1) constitutively expressed^15^. However, recent structural analysis has revealed that specifically in VRN5 there is a close packing of the central PHD superdomain and FNIII domain and this mediates its interaction with the PRC2 core complex. In contrast, VIN3 has a more open conformation of these domains and depends on VRN5 to interact with PRC2^14^.

**Figure 1:**
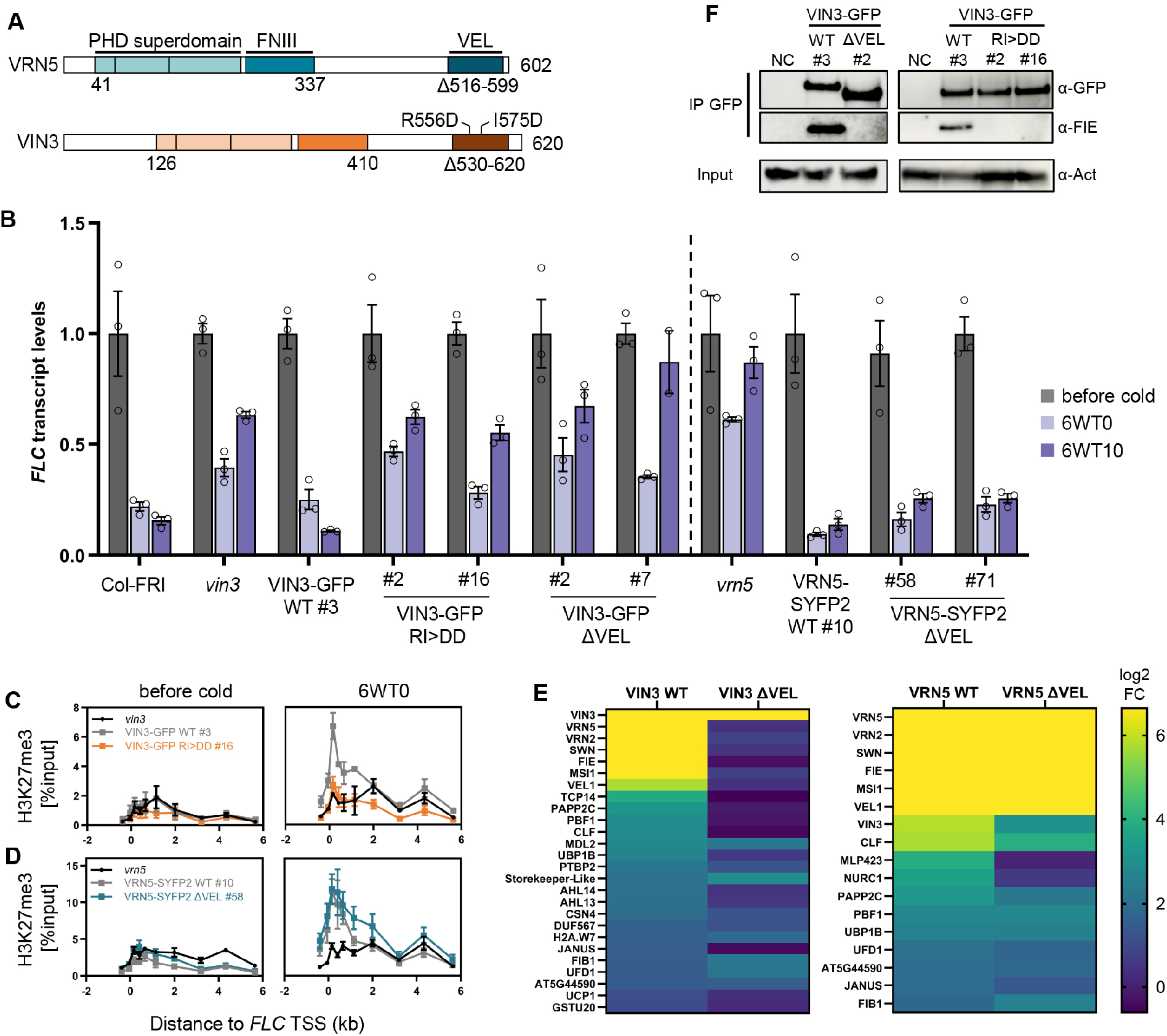
VIN3 and VRN5 VEL domain mutant lines have contrasting phenotypes. (A) Domain architecture of VEL proteins VIN3 and VRN5. Tripartite PHD superdomain: a zinc finger, an atypical PHD domain and a four-helix bundle. Above: R556D and I575D indicate polymerization-blocking point mutations in VIN3 VEL; below: amino acids deleted in VIN3-GFP ΔVEL and VRN5-SYFP2 ΔVEL. (B) qRT-PCR assays of *FLC* transcripts during a vernalization timecourse; before cold, after 6-week cold exposure (6WT0), or 10 days post-cold (6WT10). Data are relative to the geometric mean of *UBC*/*PP2A*, normalized to *FLC* before cold. Error bars represent standard deviations (*n* = 3 biological replicates). (C, D) H3K27me3 ChIP across the *FLC* locus before cold and after 6-week cold, relative to input, in (C) VIN3 or (D) VRN5 VEL domain mutant lines. TSS: transcriptional start site. Error bars represent SEM (*n* = 3 biological replicates). (E) Heatmap from IP-MS samples showing nuclear proteins co-precipitating with VIN3 and VRN5 baits in vernalized seedlings. log2 fold-change (FC) is in comparison to non-transgenic Col-FRI (adj. p-value ≤ 0.05 for wild-type proteins, *n* = 3 biological replicates). (F) Immunoblots of α-GFP immunoprecipitates using vernalized plants with VIN3-GFP transgenes, probed with α-FIE (PRC2 core). Non-transgenic Col-FRI was used as negative control (NC). Blots shown are representative of three replicates.

The VEL domain mediates homo- and heterotypic interactions between the VEL proteins. Structural and functional analyses of the VEL domain revealed it to be a head-to-tail polymerization domain, conserved through the green lineage^26^. It is only the third head-to-tail polymerization fold described in biology to date, their discovery often hindered by the insolubility of protein polymers in solution^27^. We found that the spontaneous homopolymerization of VIN3 and VEL1 via their VEL domains drives their assembly into dynamic biomolecular condensates detected by confocal microscopy following protein expression in heterologous systems^26^. Mathematical modelling of epigenetic state switching and memory at *FLC* had predicted the requirement of additional protein memory storage elements that positively feedback to reinforce themselves; for example, molecular assemblies that are maintained in sufficiently high numbers to overcome the nucleation threshold and that may persist at the locus even through the nucleosome perturbations that occur during DNA replication^19^. Nuclear assemblies of VEL proteins increase in stoichiometry during vernalization^28^, so that VEL-dependent protein polymerization and condensate formation could provide such a mechanism. Here, we therefore experimentally tested the importance of the VEL polymerization domain in the switch to the epigenetically silenced state.

## Results

### Contrasting PRC2-related phenotypes in VIN3 and VRN5 VEL domain mutants

We had previously shown that stable Arabidopsis lines expressing VIN3-GFP with single amino acid polymerization blocking point mutations located in the head and tail of the VEL domain (R556D/I575D [RI>DD], Fig. 1A), fail to rescue impaired *FLC* silencing in the *vin3* mutant^26^. This pointed to the importance of polymerization in *FLC* silencing. To extend these findings we directly compared the effect of VIN3-GFP wild-type (WT) and RI>DD transgenes when expressed at endogenous levels. Two homozygous single insertion lines with protein levels equal to a single-insertion VIN3-GFP WT line were selected from a larger T2 generation, all displaying the same non-complementation phenotype (Fig. S1). These VIN3-GFP RI>DD lines mirrored the impaired *FLC* shutdown during cold and the post-cold *FLC* reactivation observed for lines with a deletion of the entire VEL domain in VIN3-GFP (Fig. 1B and Fig. S1). Besides, H3K27me3 fails to accumulate at the *FLC* nucleation region in VIN3-GFP RI>DD (Fig. 1C). Because the RI>DD mutation disrupts VIN3 polymerization with minimal impact on the rest of the protein, we conclude that the polymerization of the VIN3 VEL domain is required to promote PRC2-mediated deposition of H3K27me3 in the *FLC* nucleation region.

Our prediction from the heterotypic interactions observed between VIN3 and VRN5 was that the VEL domain would be required for silencing by mediating the VRN5-dependent interaction between VIN3 and PRC2^14,15^. However, independent homozygous stably transformed Arabidopsis VRN5-SYFP2 ΔVEL lines with single transgene insertions in a *vrn5* mutant background (Fig. 1A and Fig. S2) showed an unexpected result; *FLC* was fully silenced in VRN5-SYFP2 ΔVEL like in a *vrn5* rescue line expressing VRN5-SYFP2 WT (Fig. 1B). In agreement, the deposition of H3K27me3 at the *FLC* nucleation region was not impaired in VRN5 ΔVEL plants (Fig. 1D).

To further understand the different phenotypes observed for the VEL domain deletions of VIN3 and VRN5 *in vivo*, we determined the interaction partners of VIN3 and VRN5 dependent on the VEL domain. Native GFP co-immunoprecipitation (coIP) followed by mass spectrometry was undertaken with respective WT and ΔVEL seedlings vernalized for six weeks. Both VIN3 and VRN5 WT proteins co-precipitated all components of one of the Arabidopsis PRC2 core complexes (VRN2, FIE, MSI1, SWN and to a lesser extent CLF) as well as each other and VEL1 as expected (Fig. 1E). The deletion of the VIN3 VEL domain resulted in the loss of all PRC2 core subunits and VRN5/VEL1. Probing for the PRC2 subunit FIE with immunoblot analysis after coIP in extracts of vernalized plants confirmed the loss of FIE interaction in both VIN3 ΔVEL and VIN3 RI>DD lines (Fig. 1F). In contrast, the interaction with PRC2 was maintained in VRN5 ΔVEL plants (Fig. 1E). This is in accordance with our previous results from heterologous coIPs in mammalian cells which mapped the PHD superdomain and the FNIII of VRN5, but not the VEL domain to the interface for PRC2 interaction^14^. Surprisingly, the interaction of VRN5 ΔVEL with VEL1 was also maintained, whereas the co-enrichment of VIN3 was strongly reduced. Although VEL1 has the same open conformation of the PHD superdomain and the FNIII domain as VIN3, it does interact specifically with the PRC2 core subunit MSI1 and this may explain the association with VRN5 ΔVEL^14^. It remains unclear whether VEL1 function might also contribute to the rescue of *FLC* silencing observed in VRN5 ΔVEL lines.

### Role of VEL domain in nuclear assembly formation of VIN3 and VRN5

The protein assemblies predicted to be involved in Polycomb epigenetic switching could be realised by VEL polymerization-based assemblies or by multimeric assemblies mediated by other interactors such as PRC2 (or combinations thereof). We therefore used the stable Arabidopsis transgenic VIN3-GFP and VRN5-SYFP2 lines to investigate the effects of VEL domain disruptions on the stoichiometry of both VIN3 and VRN5 in molecular assemblies *in vivo*. We performed single-particle tracking of the fluorescently tagged proteins with SlimVar microscopy, which uses an oblique illumination for rapid, enhanced imaging contrast enabling single-molecule detection sensitivity in intact root tips^28^. Stepwise photobleaching is employed as a calibration to determine the number of tagged molecules in detected fluorescent particles^29^.

In roots of seedlings vernalized for six weeks, we observed that the VIN3-GFP RI>DD mutations resulted in VIN3 assemblies with lower number of molecules, only 29% of tracks corresponded to assemblies larger than 10 molecules compared to 46% in VIN3-GFP WT (Fig. 2A). For VRN5-SYFP2 ΔVEL, we observed an increase in stoichiometry in comparison to the WT in seedlings vernalized for six weeks (Fig. 2B, for other time points see Fig. S3), in contrast to the decrease expected if the assemblies were VEL-mediated. Based on the track stoichiometry distribution, we calculated the periodicity, *i. e*. the intervals between nearest-neighbour peaks, of VIN3 and VRN5 within the assemblies as a proxy for their internal order. VIN3-GFP WT and VRN5-SYFP2 WT consistently showed dimeric units, which were disrupted in VIN3-GFP RI>DD (Fig. 2C) but generally maintained for VRN5-SYFP2 ΔVEL (Fig. 2D). Notably, we have previously observed VIN3 VEL domain crystals composed of protofilaments with a dimeric unit; the result of a mutual domain swapping of VEL domain helices between two VIN3 monomers^26^. It remains to be determined whether PRC2 might also contribute to VIN3 assembly formation during vernalization. Given the direct interaction between VRN5 and PRC2, we speculate that VEL-independent dimeric periodicity of VRN5 may be a result of the reported ability of PRC2 to undergo dimer formation^30-32^.

**Figure 2:**
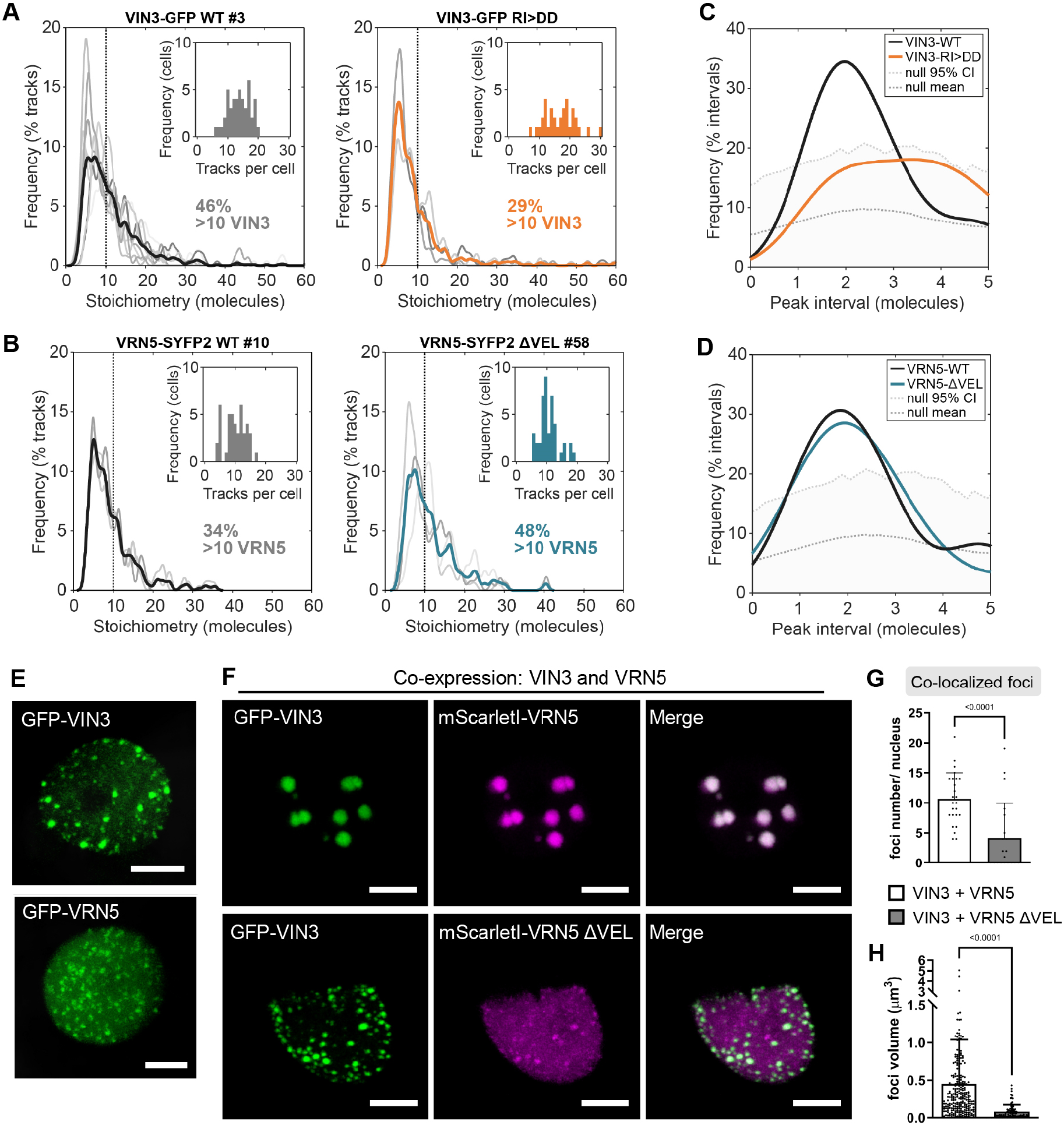
VEL-dependent VIN3 and VRN5 assemblies in vivo and after transient heterologous expression. (A, B) Stoichiometry (molecule number) distributions of SlimVar tracked assemblies of (A) VIN3-GFP WT and VIN3-GFP RI>DD and (B) VRN5-SYFP2 WT and VRN5-SYFP2 ΔVEL in root nuclei of seedlings vernalized for 6 weeks. Individual replicates (*n* = 3-8 experiments, with 9-18 nuclei each) in grey, coloured line indicates mean distribution. Insets show the frequency of tracks per cell, data collection statistics: Fig. S3. (C, D) Distributions of intervals between nearest neighbour stoichiometry peaks of tracked assemblies in (A, B). Upper dotted line indicates the null 95% confidence interval determined from simulations of random aperiodic stoichiometry, distributions which fall below are consistent with the null hypthesis^28^. (E, F) Representative confocal images of epidermal leaf cell nuclei in *N. benthamiana*, (E) transiently expressing GFP-VIN3 or GFP-VRN5, and (F) transiently co-expressing GFP-VIN3 (green) and mScarlet-VRN5 (magenta) wild-type or mutant as indicated; scale bars 5 μm. (G, H) Quantification of number per nucleus (G) and volume of foci (H) observed in (F). Error bars represent standard deviation (*n* = 20-25 nuclei for (G) and *n* = 266 (VIN3 + VRN5) or 83 (VIN3 + VRN5 ΔVEL) foci for (H)).

Because we observed no reduction in the size of VRN5 assemblies and no effect on *FLC* silencing in stable Arabidopsis lines carrying VRN5 ΔVEL, we questioned the properties of the VRN5 VEL domain. VIN3 polymerization results in the formation of biomolecular condensates^26^, so we investigated VRN5 condensate formation after transient overexpression in leaf epidermal cells of *Nicotiana benthamiana*. In comparison to the VEL- and polymerization dependent discrete condensates formed by GFP-VIN3^26^, GFP-VRN5 formed much smaller condensates, with more VRN5 protein diffusely distributed throughout the nucleus (Fig. 2E and Fig. S4A, B). Upon expression in HeLa cells, GFP-VRN5 previously also appeared diffuse^26^. Tagging VRN5 with mScarlet gave the same result as observed for GFP-VRN5 in the *N. benthamiana* system, where the deletion of the VRN5 VEL domain abolished the formation of any discrete condensates (Fig. S4C-D). These observations are consistent with VRN5 having a lower propensity to concentrate into condensates in a VEL-dependent manner. In agreement with our *in vivo* coIP results, we observed the recruitment of mScarletI-VRN5 into GFP-VIN3 to form large condensates in the *N. benthamiana* system, a co-localization that was dependent on the VRN5 VEL domain (Fig. 2F-H). Thus, the VRN5 VEL domain has some ability to homo- and heteropolymerize but its properties appear different to the VEL domain of VIN3.

### VRN5 VEL domain is not functionally equivalent to VIN3 VEL

To further examine the properties of the VRN5 VEL domain, we tested whether VRN5 VEL can functionally replace VIN3 VEL in *FLC* silencing. A VIN3-GFP construct, in which the VIN3 VEL domain was replaced with the VRN5 VEL domain (Fig. 3A), was transformed into the *vin3* mutant. *FLC* silencing during vernalization remained impaired in two independent homozygous lines as well as in multiple other lines tested in the T2 generation (Fig. 3B and Fig. S5). This suggests that VRN5 VEL is not functionally equivalent to VIN3. With equal VIN3 protein expression, coIP of the PRC2 subunit FIE was nearly undetectable in the VIN3-GFP VRN5VEL line compared to VIN3-GFP WT (Fig. 3C), most likely caused by inefficient interaction with endogenous VRN5. Equally, the heterologous co-expression of GFP-VIN3 VRN5VEL with mScarlet-VRN5 in these cells did not result in the formation of the large co-localized condensates that are observed upon co-expression of the wildtype proteins (Fig. 3D, S6). The experimentally determined structures of VIN3 and VEL1 VEL domains align closely with the Alphafold (AF) structure predictions^26^ and comparing the latter to the AF prediction for the VRN5 VEL domain again shows a close superimposition (Fig. 3E). Thus, no major structural differences are predicted to underpin the functional differences between the VEL domains. The amino acid residue I575VIN3/I664VEL1 in the VEL head interface engages in hydrophobic or electrostatic interactions with two basic residues in the complementary tail surface to mediate polymerization, blocked by the mutation I575D. A threonine in the corresponding position of Arabidopsis VRN5 and other VRN5 orthologs throughout angiosperm plants (Fig. 3F) may be functionally significant as this is a polar hydrophilic rather than a hydrophobic amino acid^26^. To investigate this, we generated the I575T mutation in recombinant VIN3VEL, bearing a lipoyl solubility tag and purified following expression in *Escherichia coli (*Fig. S7), to conduct size-exclusion chromatography coupled with multiangle light scattering (SEC-MALS). In comparison to WT VIN3VEL, polymerization was attenuated by I575T but not blocked (Fig. 3G, see comparison to the mutation I575D). This result suggests that specific amino acid differences between VIN3 and VRN5 VEL interfaces contribute to different polymerization properties, consistent with reduced VRN5 condensate formation in *N. benthamiana* cells (Fig. 2E).

**Figure 3:**
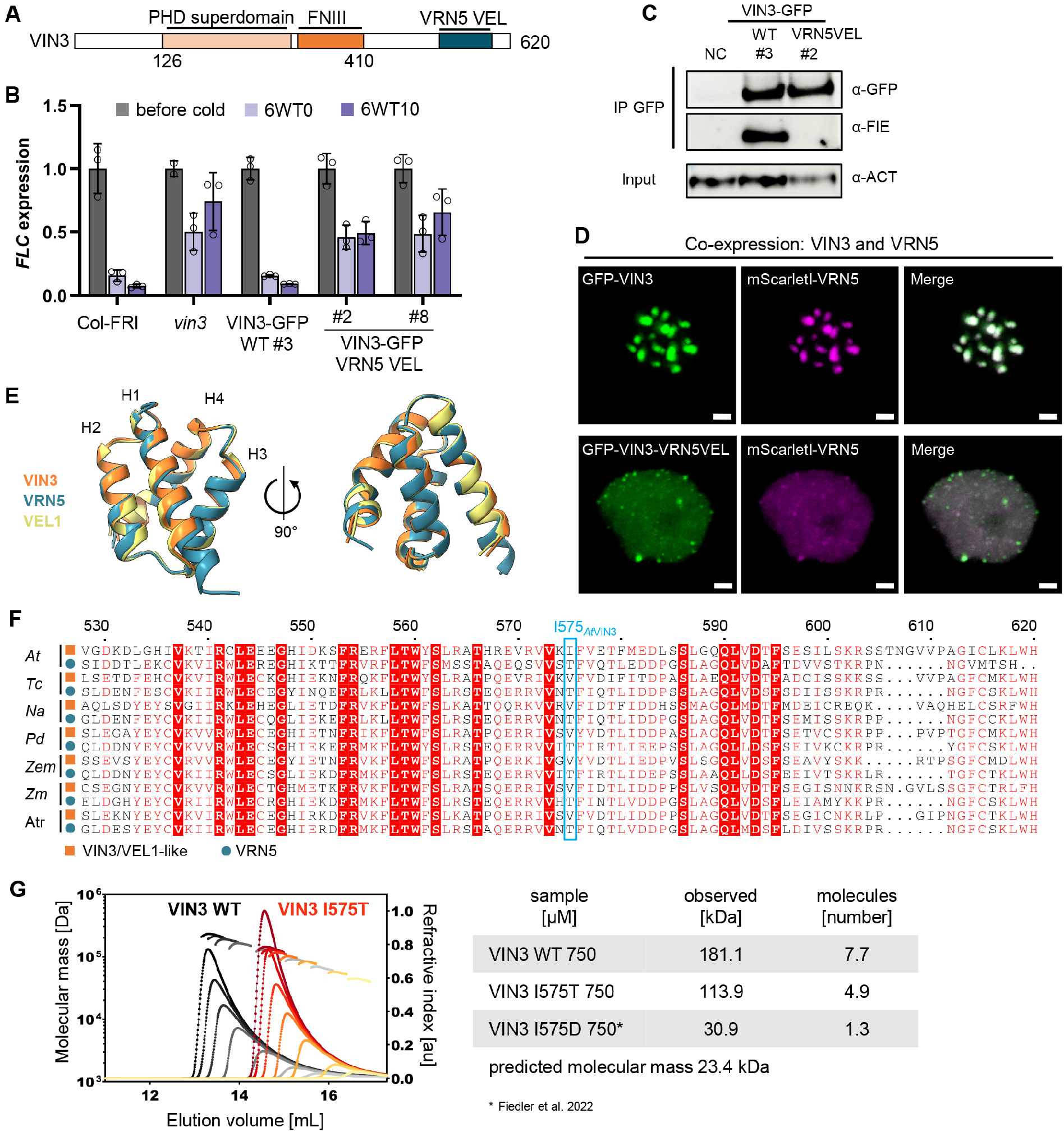
VRN5 VEL cannot replace VIN3 VEL function in planta. (A) VIN3-GFP VRN5 VEL chimera construct transformed into the *vin3* mutant. (B) qRT-PCR assays of *FLC* transcripts during a vernalization timecourse; before cold, after 6-week cold exposure (6WT0), or 10 days post-cold (6WT10). Data are relative to the geometric mean of *UBC*/*PP2A*, normalized to *FLC* before cold. Error bars represent standard deviations (*n* = 3 biological replicates). (C) Immunoblots of α-GFP immunoprecipitates from vernalized plants with indicated VIN3-GFP transgenes, probed with α-FIE (PRC2 core). Non-transgenic Col-FRI was used as negative control (NC). Blots shown are representative of three replicates. (D) Representative confocal images of epidermal leaf cell nuclei in *N. benthamiana*, transiently co-expressing GFP-VIN3 or GFP-VIN3 VRN5 VEL (green) and mScarlet-VRN5 (magenta); scale bars 2 μm. (E) Superpositions of VEL domains of Arabidopsis VRN5_515-592_ (teal), VIN3_529-601_ (orange) and VEL1_618-690_ (yellow) as predicted by Alphafold. VIN3 and VEL1 AF predictions superpose closely with experimentally determined structures^26^. (F) SEC-MALS of purified WT (grey to black) or I575T mutant (yellow to red) Lip-VIN3_VEL_ (residues 500–603) at increasing concentrations from right (50 µM) to left (1250 µM); curves: elution profiles (void volume of column at 8 mL); line traces: molar masses as derived from MALS; these are specified in the neighbouring table and also indicate numbers of molecules per oligomer at a concentration of 750 µM (note that data for VIN3 I575D is reproduced from^26^). (G) Amino acid sequence conservation of VEL domains of VRN5 (teal, defined by DLNxxxVPDLN motif in the linker region between the FNIII and VEL domains^14^) and VIN3/VEL1 orthologs (orange) throughout the angiosperm lineage. Blue borders highlight the amino acids at the position corresponding to I575_AtVIN3_. *At*: *Arabidopsis thaliana, Tc*: *Theobroma cacao, Na*: *Nicotiana attenuata, Pd*: *Phoenix dactylifera, Zem*: *Zea mays, Zm*: *Zostera marina, Atr*: *Amborella trichopoda*.

### VEL polymerization promotes multivalent VIN3 chromatin association independent of PRC2

To understand the specific contribution of VIN3 polymerization to *FLC* silencing, we determined whether the loss of interaction between VIN3 and PRC2 observed in the stable VIN3-GFP RI>DD, VIN3-GFP ΔVEL, VIN3-GFP *vrn5* as well as VIN3-GFP VRN5 VEL lines (Fig. 1E-F and Fig. 3C) would affect the association of VIN3 with the *FLC* locus. ChIP-qPCR experiments with vernalized seedlings revealed that VIN3-GFP *vrn5* and VIN3-GFP VRN5VEL showed equally high enrichment at *FLC* as VIN3-GFP WT (Fig. 4A and B), suggesting that the interaction between VIN3 and PRC2 is not *per se* required for VIN3 chromatin binding. In contrast, we observed a strongly reduced association of VIN3-GFP RI>DD and VIN3-GFP ΔVEL with the *FLC* nucleation region (Fig. 4C and S8A). The chromatin association of VRN5 ΔVEL at the *FLC* locus was WT-like in comparison (Fig. S8B). We also tested other VIN3 targets, previously identified by ChIP-seq experiments^14^, and found reduced VIN3-GFP RI>DD association at several of these loci (Fig. S9). This implicates VEL-mediated polymerization in promoting and maintaining VIN3 chromatin association with *FLC* and other loci in a PRC2-independent manner.

**Figure 4:**
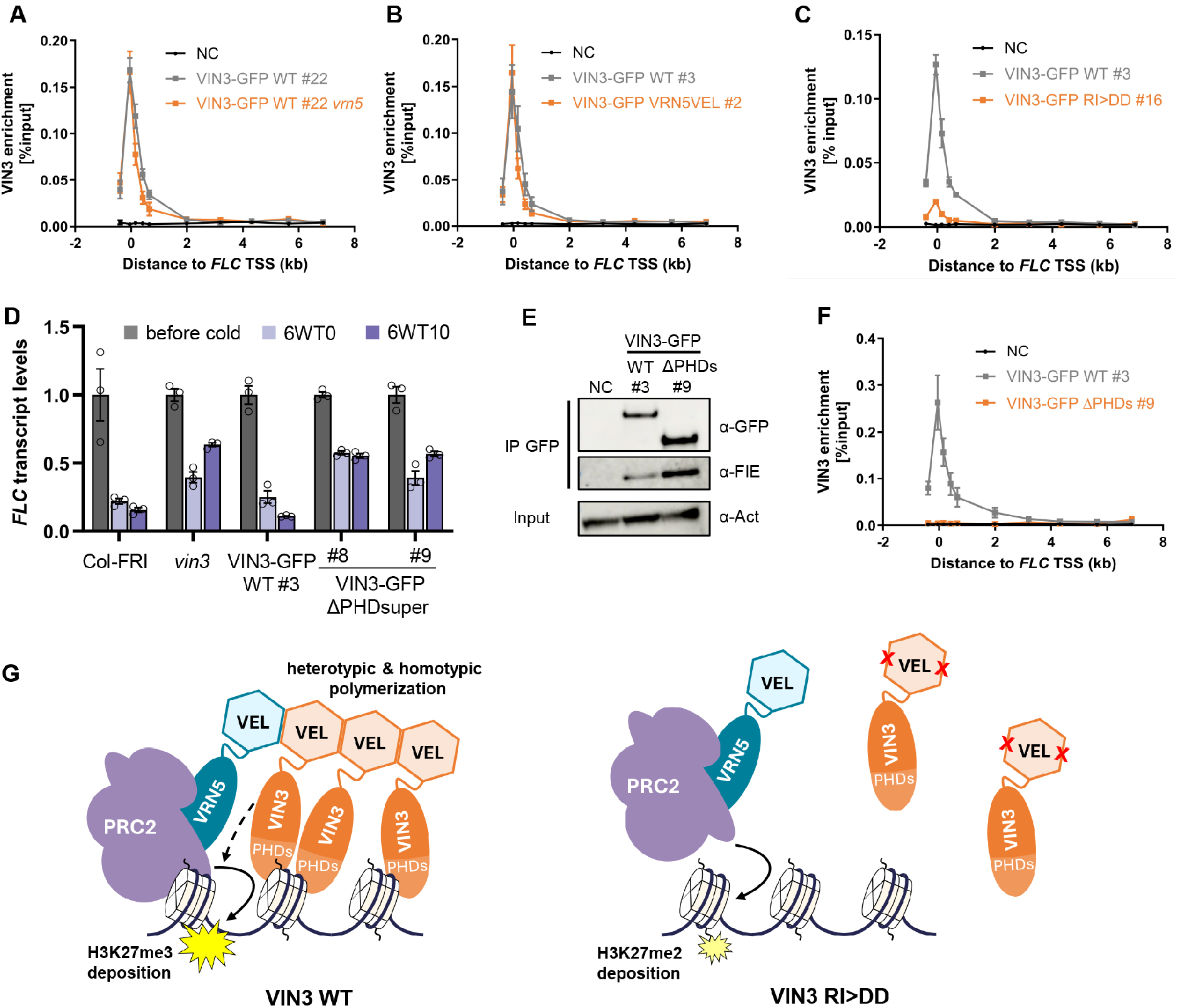
VIN3 chromatin association is promoted by VEL-mediated polymerization and chromatin binding of the PHD superdomain. (A-C, F) ChIP-qPCR showing enrichment of VIN3-GFP (wildtype or mutant as indicated) across the *FLC* locus in seedlings vernalized for 6 weeks. Non-transgenic Col-*FRI* plants were used as negative control sample (NC). Data are relative to input control, error bars represent SEM (*n* = 2-4). (D) qRT-PCR assays of *FLC* transcripts during a vernalization timecourse; before cold, after 6-week cold exposure (6WT0), or 10 days post-cold (6WT10). Data are relative to the geometric mean of *UBC*/*PP2A*, normalized to *FLC* before cold. Error bars represent standard deviations (*n* = 3 biological replicates). (E) Immunoblots of α-GFP immunoprecipitates from extracts of vernalized plants bearing the indicated VIN3-GFP transgenes, probed with α-FIE (PRC2 core subunit) antibody. Non-transgenic Col-FRI was used as a negative control (NC). Blots shown are a representative of three replicates. (G) Schematic model of VIN3 and VRN5 function in PRC2 silencing. Polymerization via the VEL domain promotes VIN3 chromatin association, mediated by emergent multivalent interactions between VIN3 PHD superdomains and chromatin ligands. This facilitates recruitment of VRN5-PRC2 via heterotypic VEL interaction but also promotes H3K27me3 nucleation by other means (dashed arrow, see discussion). Note that VIN3/VRN5 stoichiometry drawn here is a representation of the observed differences in VEL domain properties but does not correspond directly to molecule number *in vivo*.

An emerging paradigm for the function of head-to-tail protein polymerization is the increase in local concentration of polymerizing proteins and their ligand binding sites, which enhances their binding avidity for low-affinity ligands (functional affinity) ^27^. Because the head and tail interfaces of the VEL domain facilitates homo- and heterotypic interactions between the VEL proteins, both types of interactions can theoretically contribute to promote VIN3 chromatin association in such a mechanism. We observed that VIN3-GFP VRN5VEL does not restore the VRN5-dependent interaction with PRC2, yet it still binds to *FLC* efficiently; thus the properties of VIN3-VRN5VEL are sufficient for some but not all outputs achieved by the VIN3 WT protein and its VEL domain. This highlights the complexities at play *in vivo*.

We decided to test whether a functionally relevant binding site - whose avidity for its chromatin ligand might be enhanced by the VEL-dependent polymerization - is present in the VIN3 protein itself. We turned to the tripartite PHD superdomain of VIN3: although this domain is an atypical PHD domain in that it exhibits no histone H3 tail binding activity, it has a weak affinity for negatively charged DNA or RNA polymers *in vitro*^*25*^. We generated stable transgenic Arabidopsis plants carrying VIN3-GFP with a deletion of the entire PHD superdomain (ΔPHDsuper) in the *vin3* mutant background. The construct did not complement the *vin3* mutant in *FLC* vernalization time course experiments in two homozygous lines or in multiple other lines tested in the T2 generation (Fig. 4D and Fig. S10). While VIN3 ΔPHDsuper maintained its interaction with PRC2 based on FIE immunoprecipitation in vernalized seedlings (Fig. 4E), chromatin association with the *FLC* nucleation region was abolished (Fig. 4F). This demonstrates that the VIN3 PHD superdomain is necessary for chromatin association and has chromatin or chromatin-associated ligands other than histone H3 tails. Interestingly, H2A was among the significant VIN3 interactors identified by native coIP-MS (Fig. 1E), more specifically the histone variant H2A.W which is known to associate with H3K9me2-marked heterochromatin^33^. For VIN3-GFP RI>DD, the binding peak at the *FLC* nucleation region was much smaller than for VIN3-GFP WT but not entirely abolished (Fig. 4C). This may reflect the weak chromatin affinity of VIN3 monomers, mediated by the PHD superdomain in the absence of the VIN3-mediated polymerization. We previously found an interaction between VIN3 and the transcriptional repressor VAL1, which serves as an assembly platform for co-transcriptional repressors and chromatin regulators^14,34^. VAL1 binds to two RY-motifs in the first *FLC* intron and could thus also provide a sequence-specific link to the *FLC* locus^35^. However, while transgenic Arabidopsis plants carrying a point mutation in the first RY site (*FLC*-C585T) fail to nucleate H3K27me3, they still show an accumulation of H3K27me2 specifically in the nucleation region, indicating that VAL1 is only one of the multifactorial mechanisms that ensure targeting of VEL-PRC2 to this region^14,35^. In agreement with this, we found that VIN3-GFP is still recruited to *FLC* in the C585T background (Fig. S11).

## Discussion

Polymerization is one example of mechanisms that can achieve combinatorial protein inputs to promote the interaction between otherwise weakly interacting molecular components. Our findings suggest that polymerization via the VEL domain results in a high local concentration of VIN3, enhancing its avidity for chromatin by emergent multivalent interactions between VIN3 PHD superdomains and chromatin ligands (Fig. 4G). Indeed, combinatorial protein inputs have emerged to be crucial to orchestrate transcriptional output in eukaryotic gene regulation as a more general theme^36-38^, capable of buffering against noisy signals and enhancing target specificity. Experiments with transcription factors (TFs) in yeast serve as an example of the physiological relevance of such mechanisms: synthetic TFs with high affinity to *cis* regulatory motifs imposed a fitness burden that was relieved by weakly binding TFs cooperating in assemblies^39^.

Here, prolonged chromatin association of VIN3 would also increase PRC2 residence time, bridged by VRN5, to facilitate H3K27me3 nucleation (Fig. 4G). This is consistent with our previous observation that the *vin3* mutant, unlike *vrn5*, accumulates the precursor mark H3K27me2 in the nucleation region, which indicated that VIN3 is required to overcome the threshold from di-to trimethylation at *FLC*^*14*^. Similarly, the accumulation of H3K27me2 at PRC2 target sites was observed in mammalian cell lines combining knockouts of different PRC2 accessory proteins^40^. That the VRN5 VEL domain is not essential for H3K27me3 nucleation suggests that the association of the VIN3 polymer promotes H3K27me3 nucleation by other means in addition to the direct recruitment of VRN5 and PRC2, eg. by modulating chromatin properties such as nucleosome dynamics or by binding other protein effectors.

Overall, our findings for the VEL proteins have striking resemblance to a polymerization network involving multiple Polycomb Repressive Complex 1 (PRC1) subunits that engage in a combination of heterotypic and homotypic interactions to promote transcriptional repression in Drosophila. These interactions are mediated by the head-to-tail co-polymerization of SAM domains of Polyhomeotic (Ph) and Sex combs of midleg (Scm) and are linked to the Pho-repressive complex (PhoRC) subunit Sfmbt, which undergoes heterotypic SAM interactions with Scm but is unable to homopolymerize^41-43^. The DNA-binding activity of the PhoRC complex contributes to this polymerization-mediated hub to promote the nucleation of PRC1 complexes at target loci. Polymerization-disrupting mutations in the SAM domain of Ph do not alter Ph chromatin association at most genomic binding sites and have been predominantly linked to changes in long-range chromatin interactions^44^. While the relationship between VEL polymerization and chromatin looping is currently still unknown, *FLC* alleles have been observed to cluster during vernalization, which is impaired in *vrn2* and *vrn5* mutants^45^.

Like the SAM domain-dependent co-polymer^46^, the *in vivo* composition and dynamics of the VEL co-polymer will depend on the affinities for all possible homo- and heterotypic interactions between head and tail interfaces of VEL proteins. Specific amino acid residues in the polymerization interface contribute to this, observed here for attenuated polymerization of VIN3 when carrying I575 mutated to the threonine found at the corresponding position in the VRN5 VEL domain (Fig. 3G). This could also underlie the differences between VIN3 and VRN5 condensation in cells. The threonine residue is widely conserved throughout angiosperm VRN5 orthologs (Fig. 3F, see also Fig. S12 and Supplemental Table S1), all predicted to be direct interactors of PRC2 based on the compact conformation of their PHDsuper and FNIII domains^14^. In contrast, amino acids able to engage in hydrophobic interactions in the polymerization interface are prevalent in the corresponding position of angiosperm VIN3/VEL1 orthologs (Fig. 3F), which are predicted to have a more open conformation of their PHD super and FNIII domains unable to confer PRC2 interaction^14^. At least two homologs of VEL proteins, from each of the VRN5 and VIN3/VEL1-like subclasses, are present in these species, suggesting that the maintenance of VEL proteins with different PRC2 binding and polymerization properties throughout angiosperm evolution may be functionally important. Other amino acid differences in the polymerization interfaces, possibly associated with different post-translational modifications *in vivo*, are likely to influence polymerization behaviour to fine-tune the VEL polymerization network over evolutionary timescales and remain to be investigated in the future.

Taken together, our work defines distinct roles for VEL polymerization in the digital switch to the Polycomb silenced state and extends our mechanistic understanding of the principles not only underlying Polycomb switching but eukaryotic gene regulation generally.

## Supplementary Materials

Figs. S1 to S12

Supplemental Tables S1 and S2

## Acknowledgements

We thank Shuqin Chen and Caroline Smith for their excellent technical help. We are grateful to Gerhard Saalbach and Carlo Martin from the JIC proteomics facility for performing mass spectrometry analysis. We also thank all members of the Dean and Howard groups as well as Elsa Franco-Echevarría for discussions.

## Author contributions

MB and CD conceived the study. AS, GJY, APD, MF and MLN performed the investigation and analysed the data. AS and CD wrote the manuscript; all authors reviewed and edited the manuscript. MCL and CD supervised the study, acquired the funding, and were the project administrators.

## Data and materials availability

The mass spectrometry proteomics data have been deposited to the ProteomeXchange Consortium via the PRIDE partner repository with the dataset identifier PXD048844 and doi 10.6019/PXD048844. SlimVar microscopy data has been deposited to the BioStudies repository under the doi 10.6019/S-BIAD1233. All other microscopy data has been deposited to BioStudies under the doi 10.6019/S-BIAD1249.

## Materials and methods

### Plant materials and growth conditions

Seeds were surface sterilized with chlorine gas, sown on MS media plates without glucose (pH 5.7) and stratified at 4 °C in the dark for 2 days. For non-vernalized (before cold) conditions, seedlings were grown for 10 days in long-day conditions (16 h light, 8 h darkness, 20 °C). After this pre-growth, seedlings were vernalized for 6 weeks (8 h light, 16 h darkness, 5 °C) (6WT0). Vernalized seedlings were returned to before-cold conditions described above for 10 days (6WT10) for post-cold samples. For seed generation, seedlings were transferred to soil after vernalization and cultivated in a glasshouse with controlled 22°C 16 h day and 20°C 8 h night conditions. Arabidopsis Col-FRI is Col-0 with an introgressed active Sf2 FRIGIDA allele (FRI) and was previously described, as were the *vin3-1* FRI *(vin3)* ^*16*^ and the *vrn5-8* FRI (*vrn5*) mutants^14^. The transgenic lines pVIN3:VIN3-eGFP/*vin3-4* FRI (VIN3-GFP #22) ^35^, pVIN3:VIN3-eGFP/*vin3-4 vrn5-8* FRI (VIN3-GFP #22 *vrn5-8)* ^*14*^, pVIN3:VIN3-eGFP/*vin3-1* FRI (VIN3-GFP WT #3-8) and pVIN3:VIN3-eGFP R556D/I575D *vin3-1* FRI (VIN3-GFP RI>DD) ^26^ were described previously. The previously transcribed transgene *FLC*-C585T ^35^ was transformed into *FLClean* FRI^*47*^ and then crossed to pVIN3:VIN3-GFP #22/*vin3-4* FRI to generate VIN3-GFP *FLC*-C585T #36.

### Generation of transgenic Arabidopsis lines

All primers used for cloning newly generated constructs are listed in Supplemental Table S2. The genomic pENTR pVIN3::VIN3-GFP construct^35^ was modified to make pENTR pVIN3::VIN3-GFPΔPHDsuper and pENTR pVIN3::VIN3-GFP ΔVEL by Quickchange, using Phusion DNA Polymerase (Thermo Scientific). To generate pENTR pVIN3::VIN3-GFP VRN5VEL, the VRN5 VEL domain was amplified from pENTR pVRN5-SYFP2 and swapped into pENTR pVIN3-VIN3-GFP by restriction-free cloning.

To generate pENTR pVRN5::VRN5-SYFP2, the VRN5 genomic region including endogenous promoter and terminator were amplified from genomic Col-0 DNA and cloned into pENTR using In-Fusion cloning (Takara). SYFP2 was then inserted with In-Fusion cloning, followed by the insertion of a 10 amino acid linker between VRN5 and SYFP2 with Quickchange PCR using Phusion DNA Polymerase (Thermo Scientific). This wild-type construct was modified by Quickchange, using Phusion DNA polymerase, to generate pENTR pVRN5::VRN5-SYFP2 ΔVEL. All plasmids were verified by sequencing. The pENTR-constructs were transferred to the binary vector pSLJ-DEST (based on pSLJ755I6) with LR reaction (Invitrogen) and then transformed into *vin3-1 FRI* or *vrn5-8* FRI mutants mediated by Agrobacterium C58 using the floral dip method. Transgene copy number was determined in T1 transformants by IDna Genetics (Norwich Research Park).

### RNA extraction and qRT-PCR

RNA was extracted as described^35^, using acidic phenol followed by lithium chloride precipitation. RNA was DNase treated with Turbo DNA Free DNase, then transcribed into cDNA with SuperScript Reverse Transcriptase IV (both Life Technologies) with gene-specific reverse primers (Supplemental Table S1). qPCR was performed using SYBRGreen Master Mix II on a LightCycler 480 II (both Roche) with primer pairs listed in Supplemental Table S1.

### Co-immunoprecipitation and immunoblotting

For co-immunoprecipitation (coIP) analysis followed by immunoblot analyses, total proteins were extracted from 2-3 aliquots of 3 g frozen ground Arabidopsis seedling tissue with IP buffer (50 mM Tris-HCl pH 7.5, 150 mM NaCl, 0.5% NP-40, 1% Triton-X, EDTA-free protease inhibitor cocktail [Roche]). Lysates were cleared by filtering through miracloth followed by centrifugation (6,000 g, 30 min, 4°C) and then incubated with GFP-Trap (Chromotek) for 4h. Immunoprecipitates were washed four times with IP buffer and eluted by boiling in 4x NuPAGE LDS sample buffer for 10 min. Input and coIP fractions were separated by polyacrylamide gel electrophoresis (SDS-PAGE) and blotted onto polyvinylidine difluoride (PVDF) membranes. Primary antibodies anti-GFP (11814460001, Roche) and anti-FIE (AS12 2616, Agrisera) were diluted 1:1000, anti-Actin was diluted 1:5000 (AS132640, Agrisera). Secondary antibodies were HRP-coupled. Blots were washed with TBS containing 0.05% Tween-20 and developed with SuperSignal West Femto Maximum Sensitivity Substrate (Thermo Scientific).

### Co-immunoprecipitation and mass spectrometry

For coIP followed by mass spectrometry, total proteins were extracted from 3 g of frozen ground Arabidopsis seedling tissue with the IP buffer described above, with the addition of PhosSTOP according to manufacturer instructions (4906845001, Roche). IP was performed as described above. Immunoprecipitates were eluted by boiling for 15 min in 20 mM Tris-HCl pH 8, 2% SDS. Proteins in the eluate were precipitated with chloroform/methanol (1:4) on ice for 30 min, the pellet was then washed twice with methanol and once with acetone before drying. Protein pellets were resuspended in 50 µl of 1.5% sodium deoxycholate (SDC; Merck) in 0.2 M EPPS-buffer (Merck), pH 8.5 and vortexed under heating. Cysteine residues were reduced with dithiothreitol, alkylated with iodoacetamide, and the proteins digested with trypsin in the SDC buffer according to standard procedures. After the digest, the SDC was precipitated by adjusting to 0.2% trifluoroacetic acid (TFA), and the clear supernatant subjected to C18 SPE using home-made stage tips with C18 Reprosil_pur 120, 5 µm (Dr Maisch, Germany). Aliquots were analysed by nanoLC-MS/MS on an Orbitrap Eclipse™ Tribrid™ mass spectrometer coupled to an UltiMate^®^ 3000 RSLCnano LC system (Thermo Fisher Scientific, Hemel Hempstead, UK). The samples were loaded onto a trap cartridge (PepMap™ Neo Trap Cartridge, C18, 5um, 0.3×5mm, Thermo) with 0.1% TFA at 15 µl min^-1^ for 3 min. The trap column was then switched in-line with the analytical column (Aurora Frontier TS, 60 cm nanoflow UHPLC column, ID 75 µm, reversed phase C18, 1.7 µm, 120 Å; IonOpticks, Fitzroy, Australia) for separation at 55°C using the following gradient of solvents A (water, 0.1% formic acid) and B (80% acetonitrile, 0.1% formic acid) at a flow rate of 0.26 µl min^-1^ : 0-3 min 1% B (parallel to trapping); 3-10 min increase B (curve 4) to 8%; 10-102 min linear increase B to 48; followed by a ramp to 99% B and re-equilibration to 0% B, for a total of 140 min runtime. Mass spectrometry data were acquired with the FAIMS device set to three compensation voltages (−35V, -50V, -65V) at standard resolution for 1.0 s each with the following MS settings in positive ion mode: OT resolution 120K, profile mode, mass range m/z 300-1600, normalized AGC target 100%, max inject time 50 ms; MS2 in IT Turbo mode: quadrupole isolation window 1 Da, charge states 2-5, threshold 1e^4^, HCD CE = 30, AGC target standard, max. injection time dynamic, dynamic exclusion 1 count for 15 s with mass tolerance of ±10 ppm, one charge state per precursor only.

The mass spectrometry raw data were processed and quantified in Proteome Discoverer 3.1 (Thermo); all mentioned tools of the following workflow are nodes of the proprietary Proteome Discoverer (PD) software. The *A. thaliana* TAIR10 protein database (arabidopsis.org; 32785 entries) was modified by removing accessions AT4G30200.1, AT4G30200.3, and AT4G30200.4 corresponding to 3 versions of the VEL1 protein. Only AT4G30200.2 corresponding to the canonical version of VEL1 was left in the database for clearer search and quantification results. The database search including a decoy search was performed with Mascot Server 2.8.3 (Matrixscience, London; in house server) with a fragment tolerance of 0.5 Da, enzyme trypsin with 2 missed cleavages, variable modifications were oxidation (M), Acetyl (Protein N-term), phosphorylation (STY), methylation/dimethylation/trimethylation (K); fixed modification carbamidomethyl (C). Validation in PD was then performed using Percolator based on q-values and FDR targets 0.01 (strict) and 0.05 (relaxed). The workflow included the Minora Feature Detector with min. trace length 7, S/N 3, PSM confidence high. The consensus workflow in the PD software was used to evaluate the peptide identifications and to measure the abundances of the peptides based on the LC-peak intensities. For identification, an FDR of 0.01 was used as strict threshold, and 0.05 as relaxed threshold.

For quantification, three replicates per condition were measured. In PD3.1, the following parameters were used for ratio calculation: normalisation on total peptide abundances, protein abundance-based ratio calculation using the top3 most abundant peptides, missing values imputation by low abundance resampling, hypothesis testing by t-test (background based), adjusted p-value calculation by BH-method. The results were exported into a Microsoft Excel table including data for protein abundances, ratios, p-values, number of peptides, protein coverage, the search identification score and other important values.

Protein lists obtained for VIN3 and VRN5 wild-type proteins were filtered for interactors that were positively enriched in comparison to a Col-FRI non-transgenic control sample with an adjusted p-value ≤ 0.05. Enrichment ratios of interactors predicted to have a nuclear localization were log2-transformed for both WT and mutant samples to generate the heatmaps visualising the IP-MS results. The full mass spectrometry proteomics data have been deposited to the ProteomeXchange Consortium via the PRIDE partner repository with the dataset identifier PXD048844 and 10.6019/PXD048844.

### Chromatin immunoprecipitation (ChIP)

Histone ChIP was performed with 2 g of formaldehyde-crosslinked material as described previously^14^ with the following modifications: after nuclei extraction with Honda buffer, nuclei were layered on a Percoll density gradient (75%/40% Percoll in Honda) and extracted from the interface between these layers after centrifugation (7.000 x *g* in a swing bucket rotor for 30 min at 4 °C) prior to lysis of nuclei. Immunoprecipitation was performed with antibodies α-H3K27me3 (Abcam, ab192985) and α-H3 antibody (Abcam, ab1791), using 3 µg per IP reaction. Non-histone ChIP (VIN3/VRN5) was performed as described for GFP/YFP-tagged proteins^14^. For lines with endogenous level VIN3 expression, each ChIP replicate was generated by pooling chromatin from three aliquots of 3 g of formaldehyde-crosslinked material for IP. Immunoprecipitation was performed with α-GFP (Abcam, ab290) using 3 µg per IP reaction.

### Heterologous Nicotiana benthamiana transfections

The generation of *p35S:Ω-GFP-VIN3* was described previously^26^. This plasmid was modified with seamless megaprimer cloning to generate *p35S:Ω-GFP-VRN5* and *p35S:Ω-mScarlet-VRN5* with the coding sequence of VRN5. Plasmids were transformed into *Agrobacterium tumefaciens* GV3101 using electroporation. Agrobacteria containing the desired construct at OD600 0.05 were equally co-infiltrated with the silencing suppressor P19 into three-week-old *Nicotiana benthamiana* leaves. Confocal imaging of infiltrated epidermal leaf cells of *N. benthamiana* was performed on a Leica confocal Stellaris 8 microscope using a 63x/1.2 water objective and 4x zoom, excitation at 488 nm, detection at 507-542 nm for GFP and excitation at 561 nm, detection at 575-625 nm for mScarlet. Images were acquired 24 hr after infiltration with a laser speed of 600 Hz, a typical Z-step size of 4.7 µm and a pinhole size of 1 AU. The same settings were used at all imaging sets to allow direct comparison between constructs. The analysis was performed in Arivis Vision4D ver. 4.1.0. (Zeiss). Firstly, the blob finder algorithm was applied to the GFP channel using a diameter value of 0.8 mm, a probability threshold of 50%, and a split sensitivity of 65%. Then, the blob finder algorithm was applied to the mScarlet channel using the same settings for diameter value, probability, and split sensitivity. Afterwards, the intersection between the output of the two blob finder operations was calculated. Finally, metrics such as volume were computed for the objects generated by each of the blob finder operations, as well as for their intersection.

### SlimVar microscopy and single-assembly analysis

The SlimVar technique detects rapidly diffusing single molecular assemblies inside root tip nuclei. The microscope was employed in single-colour mode as described previously^28^. Briefly, prepared seedlings were laid on a pad of MS growth media with 1% agarose on a standard slide, then coated with filtered MS media and sealed under #1.5 coverslips.

Individual nuclei within the outer three cell layers of the meristematic region of each root tip were identified in brightfield using a 100× NA 1.49 objective and centred in a region of interest no greater than 10 µm × 16 µm (190 × 300 pixel). Each nucleus of GFP- and SYFP2-labelled lines was illuminated rapidly at 3 kW cm^-2^ at an oblique angle of 60° with a collimated 488 nm or 514 nm laser respectively, and detected with a high performance sCMOS camera (Teledyne Prime95B) through a 500-550 nm or 525-575 nm emission filter respectively. The exposure time was 10 ms per frame, at a sampling rate of ∼80 fps, with the sequence length sufficient to capture complete photobleaching down to single-molecule steps, typically ∼1000 frames. Further independent measurements were taken with at least >3 nuclei per root and >3 roots per plate, for >3 independent growth and vernalization replicates (for statistics see Fig. S3C).

In post-processing analysis, also following^28^, diffraction-limited foci were extracted from each image sequence and connected into tracks that we identified with molecular assemblies. The stoichiometry of each track was estimated based on its initial fluorescent intensity, compared to that of the single label steps during late-stage photobleaching. Stoichiometry distributions were collated from populations of tracked assemblies for each line and condition. The periodicity of each stoichiometry distribution was estimated relative to a null distribution collated from simulated populations of uniform, random stoichiometry, to detect the presence of any regular structural subunits within assemblies^28,48^. Noting the previous observation that VRN5 assemblies above a threshold size of 10 molecules have a greater tendency to colocalize at the *FLC* locus as a result of vernalization^28^, the proportion of assemblies above a stoichiometry of 10 was also determined for each population in this study.

### SEC-MALS

Recombinant 6xHisLip-tagged VIN3VEL (residues 500–603), either WT or I575T mutation were expressed and purified from *E. coli* and then used for SEC-MALS as previously described^26^, with the following modification: SEC-MALS was performed using a Superose6 Increase 10/300 column.

