## Supplementary Material for "Multifunctional polymerization domains determine the onset of epigenetic silencing in Arabidopsis"

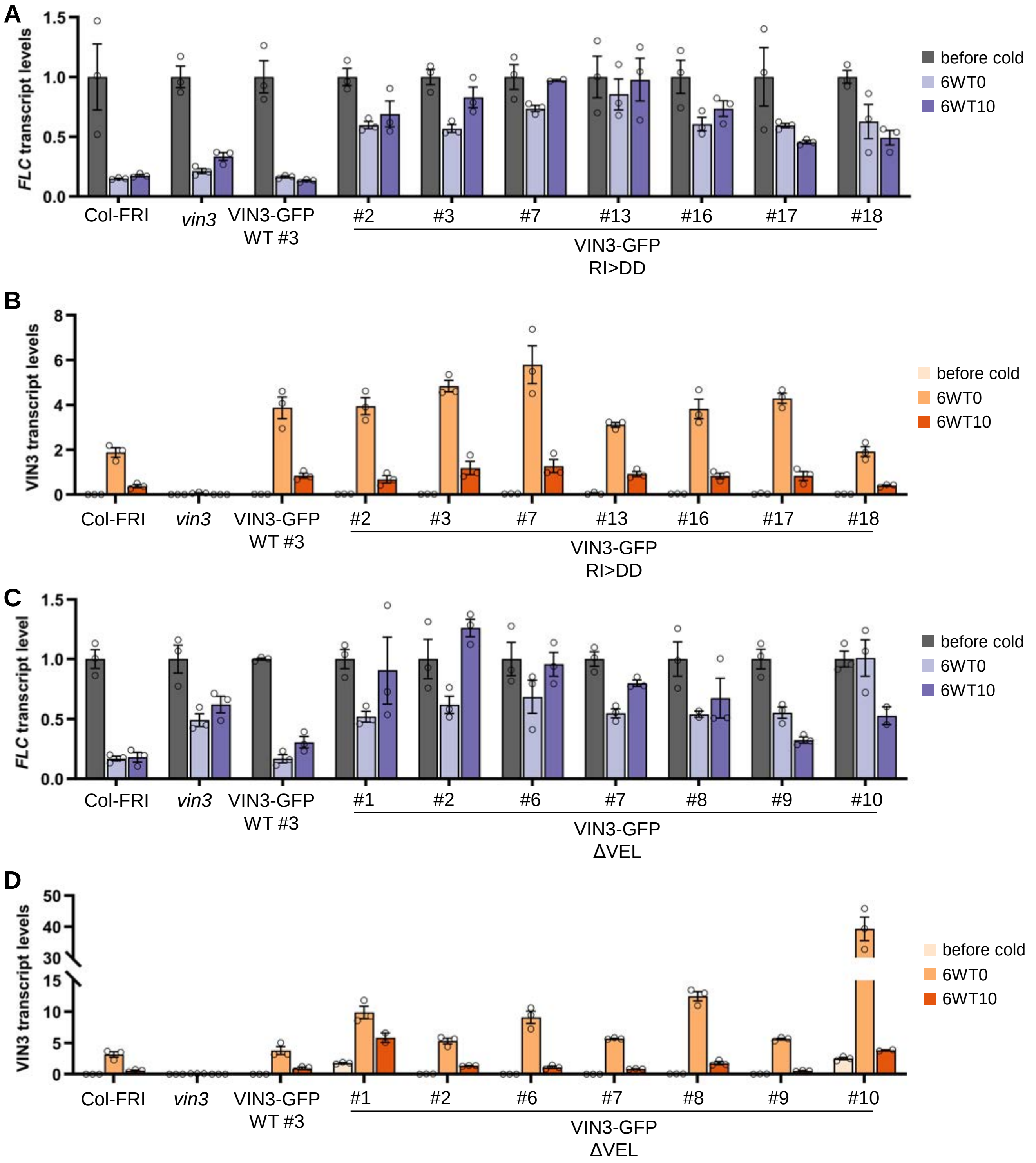


**Fig. S1: Complementation analysis of VIN3-GFP RI>DD and ΔVEL mutant lines in the *vin3* background**

(A-D): qRT-PCR assays of *FLC* (A, C) and *VIN3* (B, D) transcript levels during a vernalization timecourse. RNA was extracted from homozygous plants (Col-FRI, *vin3*, VIN3-GFP WT #3) or from seven individual segregating T2 plant lines, grown on media supplemented with the herbicide PPT to select for resistant plants, before vernalization (before cold), at the end of a 6-week cold exposure (6WT0), or 10 days post-cold (6WT10). Data presented are relative to the geometric mean of *UBC* and *PP2A*. *FLC* transcript levels are normalized to *FLC* levels before the cold. Error bars represent standard deviations ($n$ = 3 biological replicates).

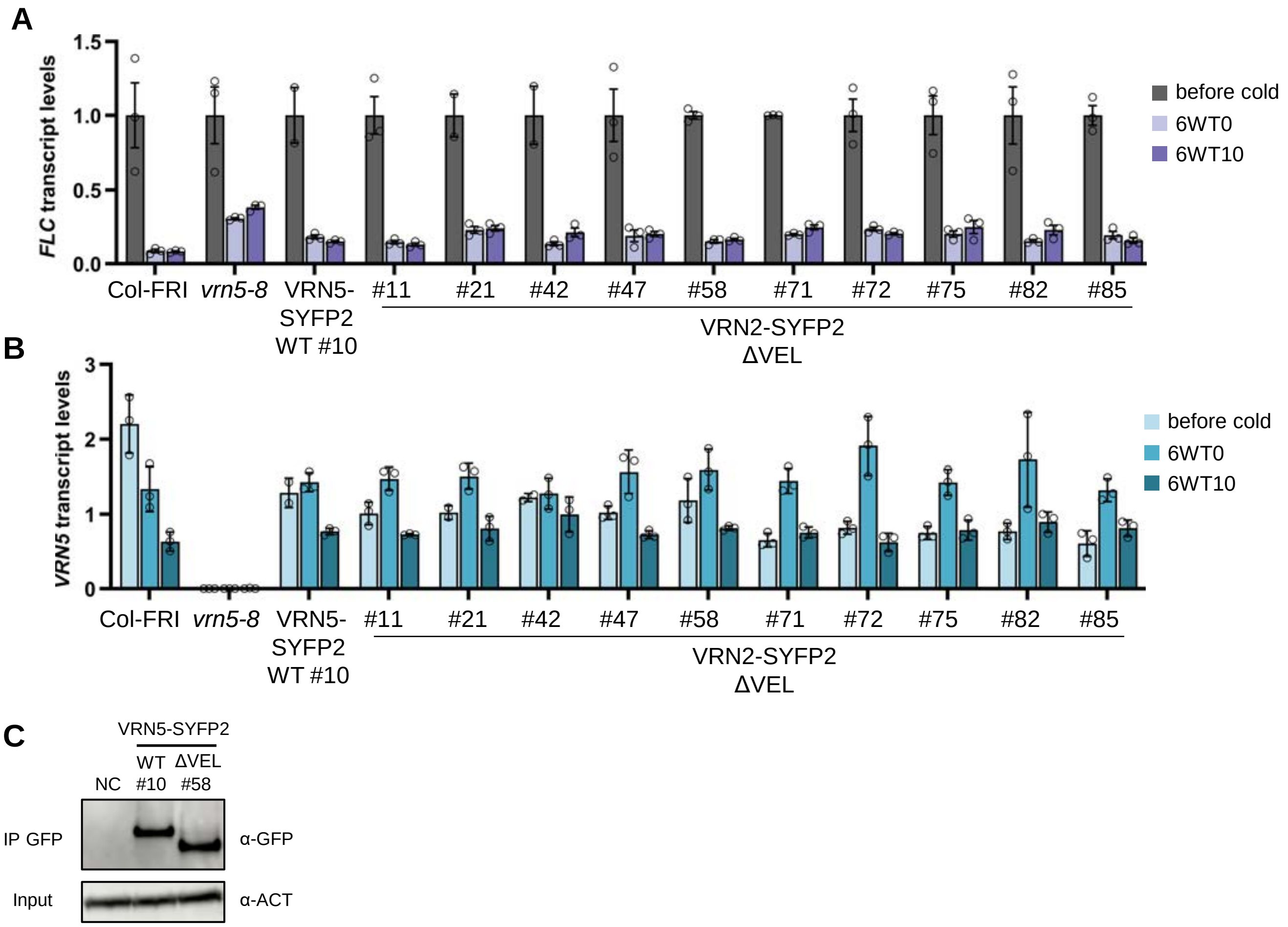


**Fig. S2: Complementation analysis of VRN5-SYFP2 WT and ΔVEL mutant lines in the *vrn5* background**

(A, B) qRT-PCR assays of *FLC* (A) and VRN5 (B) transcript levels during a vernalization timecourse. RNA was extracted from homozygous plants (Col-FRI, *vrn5*, VRN5-SYFP2 WT #10) or from 10 individual segregating T2 plant lines (all single transgene insertion), grown on media supplemented with the herbicide PPT to select for resistant plants, before vernalization (before cold), at the end of a 6-week cold exposure (6WT0), or 10 days post-cold (6WT10). Data presented are relative to the geometric mean of *UBC* and *PP2A*. *FLC* transcript levels are normalized to *FLC* levels before the cold. Error bars represent standard deviations (*n* = 3 biological replicates). (C) Immunoblots of α-GFP immunoprecipitates from extracts of vernalized plants (6 weeks) bearing the indicated VRN5-SYFP2 transgenes. Non-transgenic Col-FRI was used as a negative control (NC). Blots shown are a representative of three replicates.

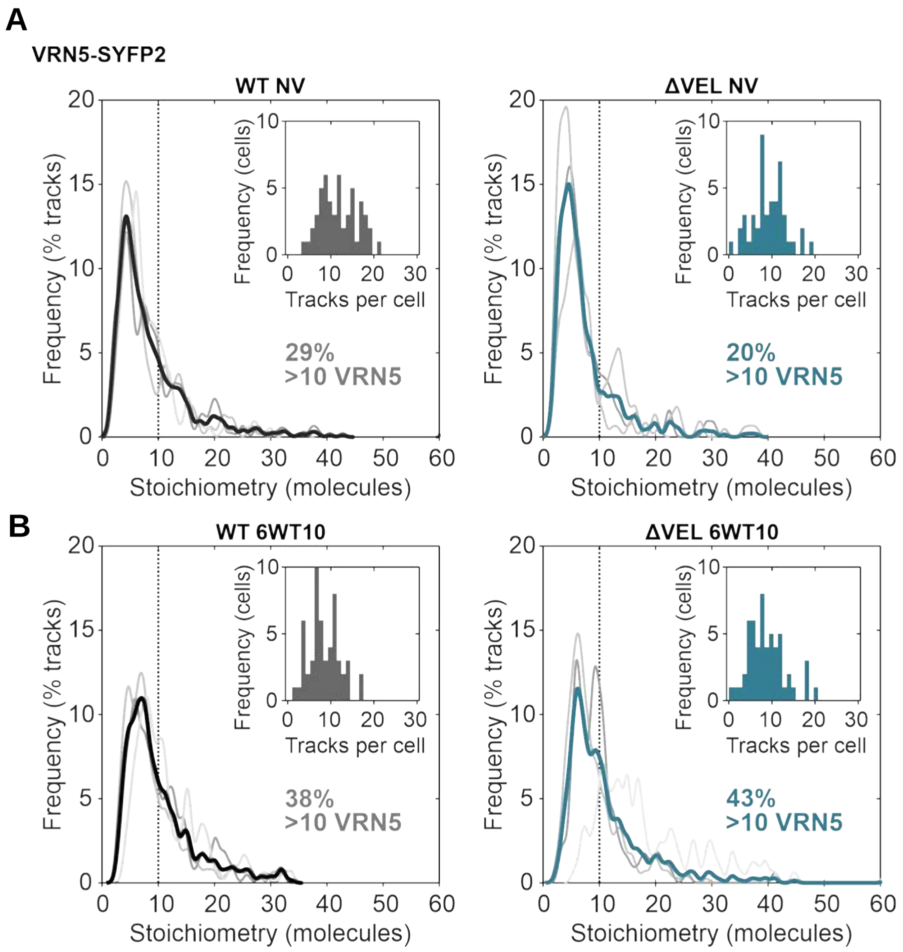


C

| Timepoint | Line | Nuclei | Tracks | Mean tracks per cell | ± std.err. | Tracks % S>10 | ± std.err. |
|---|---|---|---|---|---|---|---|
| 6WT0 | VIN3-WT | 30 | 372 | 12.4 | 0.6 | 46% | 4% |
| 6WT0 | VIN3-RIDD | 34 | 567 | 16.7 | 0.7 | 29% | 3% |
| NV | VRN5-WT | 52 | 621 | 11.9 | 0.6 | 29% | 1% |
| NV | VRN5-ΔVEL | 47 | 444 | 9.4 | 0.5 | 20% | 6% |
| 6WT0 | VRN5-WT | 37 | 386 | 10.4 | 0.5 | 34% | 4% |
| 6WT0 | VRN5-ΔVEL | 43 | 394 | 9.2 | 0.5 | 48% | 5% |
| 6WT10 | VRN5-WT | 54 | 468 | 8.7 | 0.5 | 38% | 5% |
| 6WT10 | VRN5-ΔVEL | 55 | 492 | 8.9 | 0.5 | 43% | 14% |

**Fig. S3: In vivo VEL protein assemblies at additional time points (related to figure 2)**

(A, B) Distributions of stoichiometry (molecule number) of tracked assemblies of VRN5-SYFP2 WT and VRN5-SYFP2 ΔVEL at (A) non-vernalized NV or (B) post-vernalized 6WT10 conditions, in root nuclei of seedlings determined with SlimVar. Individual replicates (n = 3 experiments, with 9-18 nuclei each) are shown in grey, the coloured line indicates the mean distribution. Insets show the frequency of tracks per cell. (C) SlimVar statistics for data shown in main figure 2 and this supplemental figure.

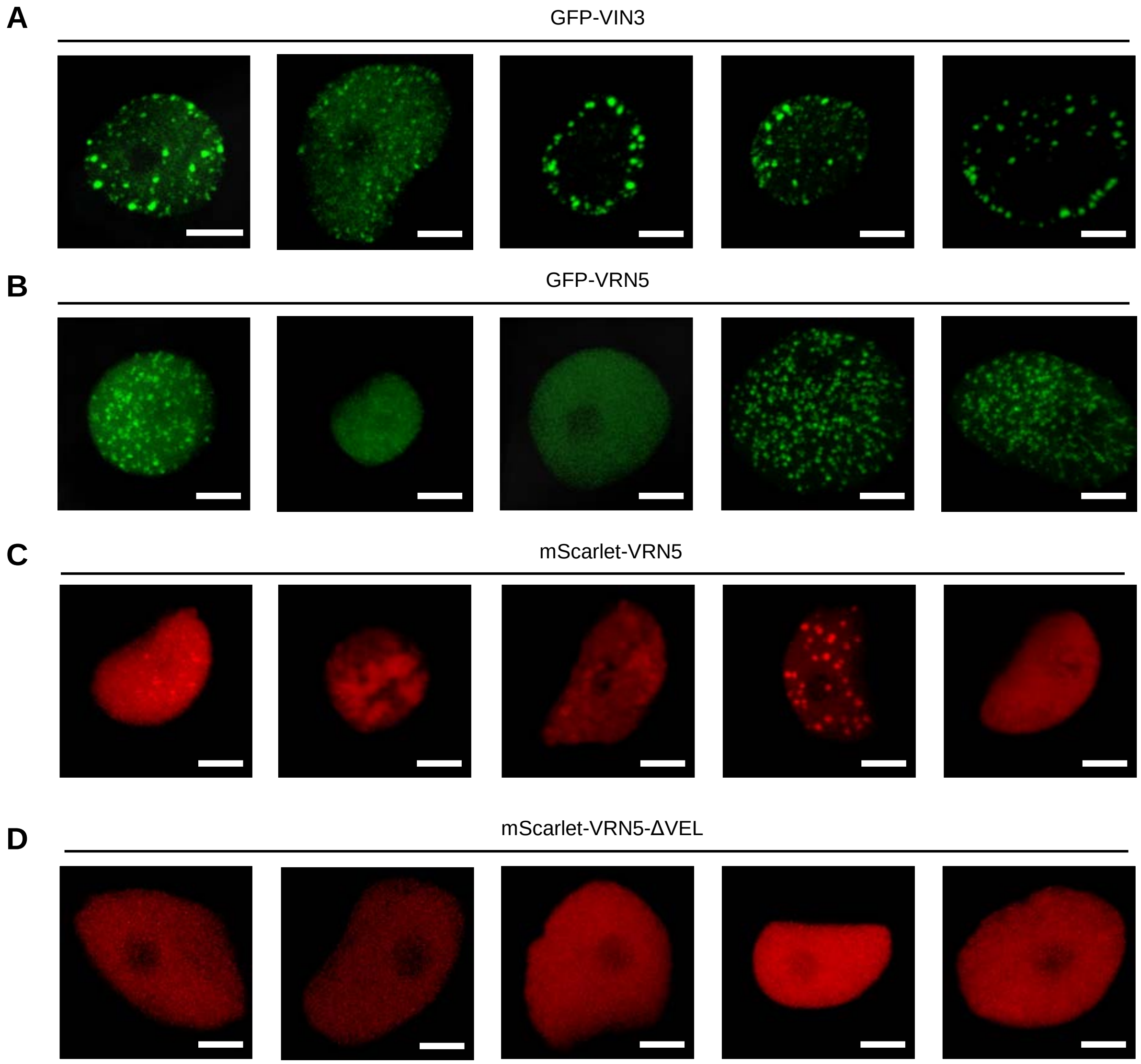


**Fig. S4: VIN3 and VRN5 foci formation upon transient expression, related to Figure 2** Additional confocal images of epidermal leaf cell nuclei in *N. benthamiana,* transiently expressing GFP-VIN3 (A), GFP-VRN5 (B), mScarlet-VRN5 (C) or mScarlet-VRN5 ΔVEL (D). For (C, D), whole image brightness was adjusted based on the mean intensity of mScarlet in the nucleus. Scale bars: 5 μm

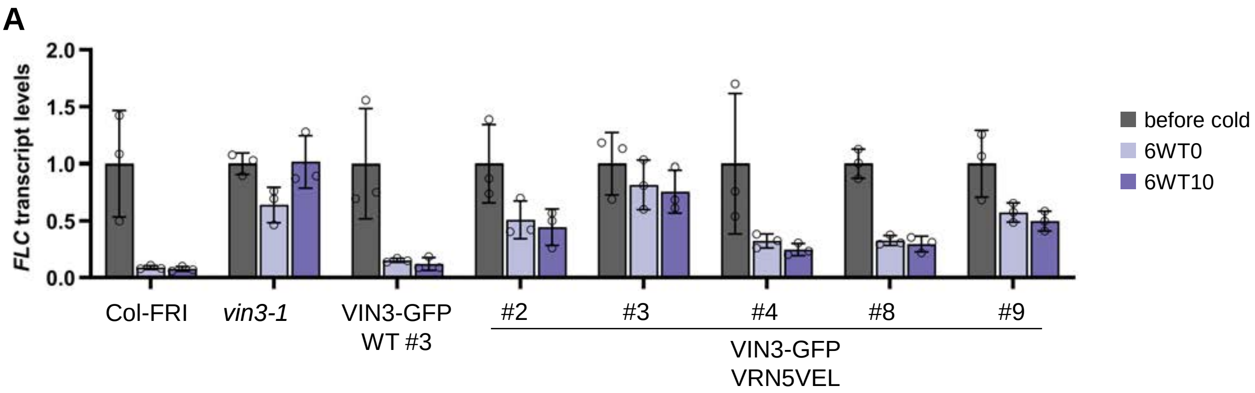


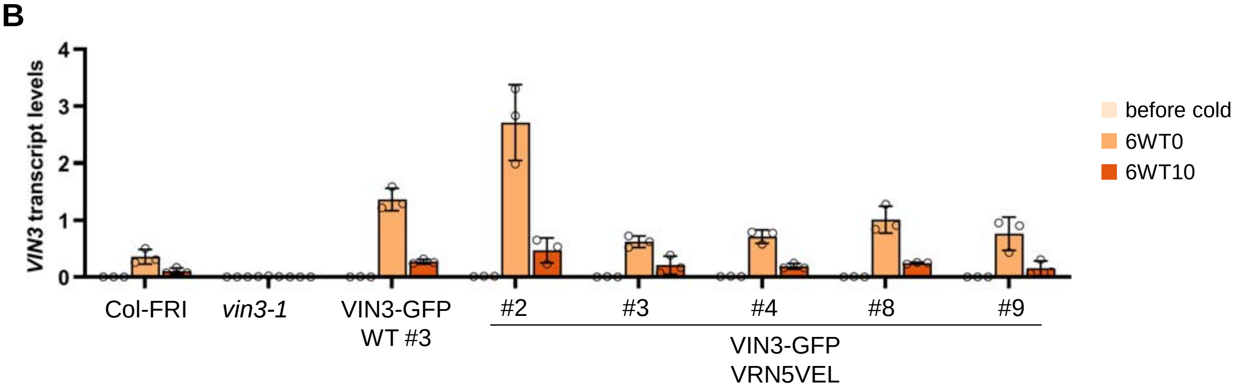


**Fig. S5: Complementation analysis of VIN3-GFP VRN5VEL mutant lines in the *vin3* background**

(A, B): qRT-PCR assays of *FLC* (A) and *VIN3* (B) transcript levels during a vernalization timecourse. RNA was extracted from homozygous plants (Col-FRI, *vin3*, VIN3-GFP WT #3) or from five individual segregating T2 plant lines, grown on media supplemented with the herbicide PPT to select for resistant plants, before vernalization (before cold), at the end of a 6-week cold exposure (6WT0), or 10 days post-cold (6WT10). Data presented are relative to the geometric mean of *UBC* and *PP2A*. *FLC* transcript levels are normalized to *FLC* levels before the cold. Error bars represent standard deviations (*n* = 3 biological replicates).

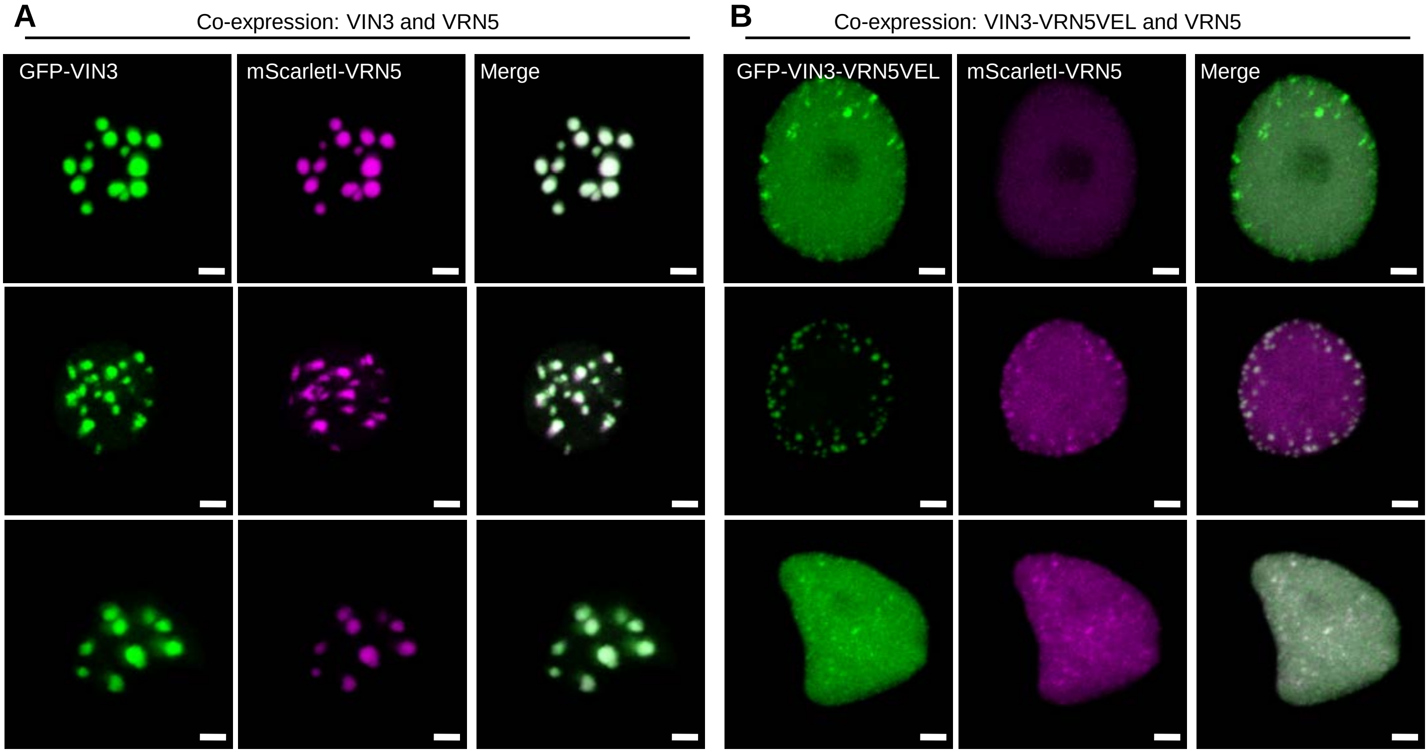


**Fig. S6: Transient co-expression of VIN3 and VRN5, related to Figure 3**

(A, B) Additional confocal images of epidermal leaf cell nuclei in *N. benthamiana,* transiently expressing GFP-VIN3 and mScarlet-VRN5 (A) or GFP-VIN3 VRN5VEL and mScarlet-VRN5 (B). Scale bars: 2 µm

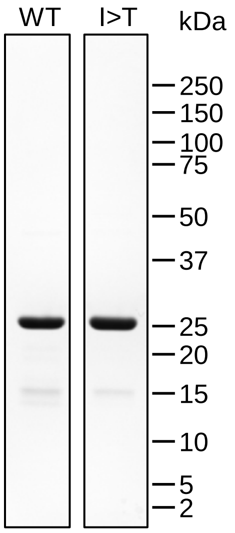


**Fig. S7: Coomassie staining of WT and mutant (I575T) purified Lip-VIN3$_{VEL}$ (related to Fig. 3G)**

Coomassie-stained protein gel, loaded with 2 ug protein per lane.

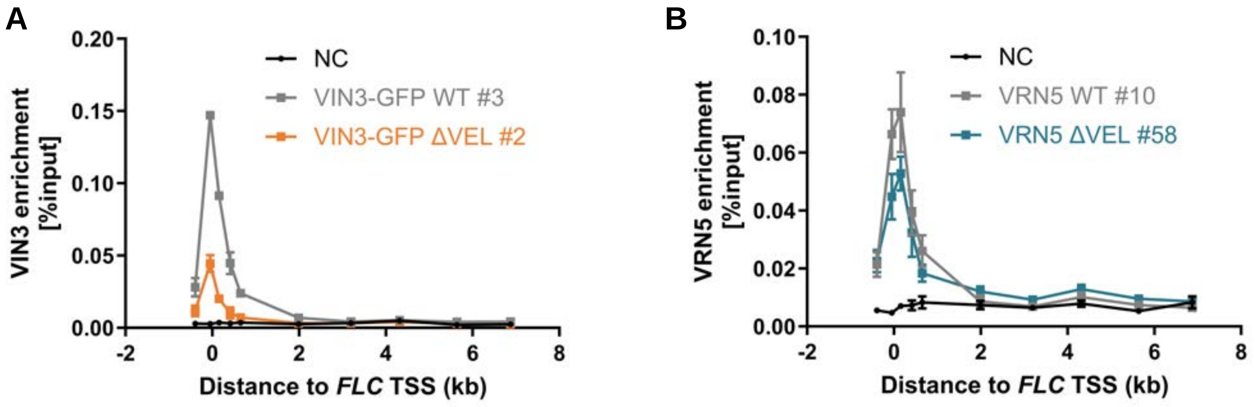


**Fig. S8: Association of VIN3-GFP ΔVEL and VRN5-SYFP2 ΔVEL with the *FLC* locus**

(A, B) ChIP-qPCR showing enrichment of VIN3-GFP (A) and VRN5-SYFP2 (B) with wildtype or mutant protein as indicated across the *FLC* locus in seedlings vernalized for 6 weeks. Non-transgenic Col-FRI plants were used as a negative control sample (NC). Data are shown relative to an input control, error bars represent SEM (*n* = 2-6).

A

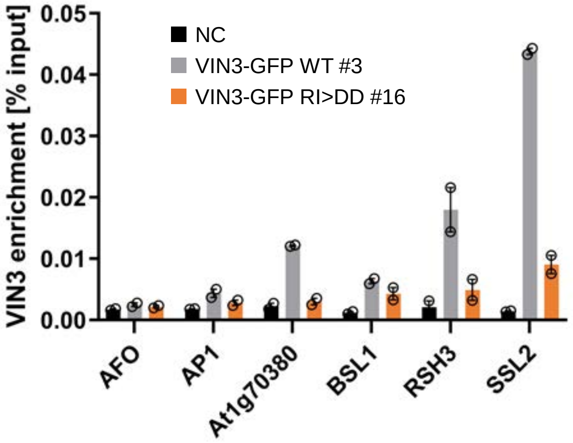


**Fig. S9: Chromatin association of VIN3-GFP RI>DD is reduced in comparison to VIN3-GFP WT at several VIN3 targets**

(A) ChIP-qPCR showing enrichment of VIN3-GFP (wildtype or mutant as indicated) at VIN3 targets genes identified by ChIP-seq in seedlings vernalized for 6 weeks. Non-transgenic Col-*FRI* plants were used as a negative control sample (NC). Data are shown relative to an input control, error bars represent SEM (*n* = 2).

A

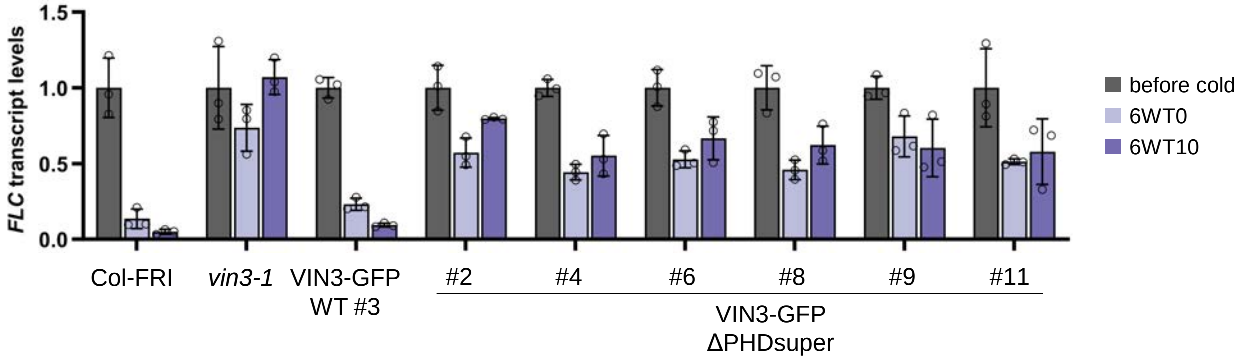


B

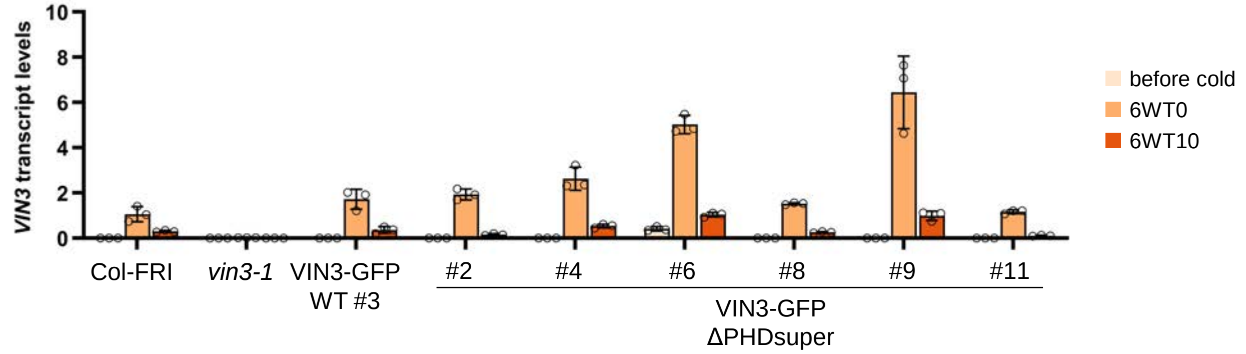


**Fig. S10 Complementation analysis of VIN3-GFP ΔPHD superdomain mutant lines in the *vin3* background**

(A): qRT-PCR assays of *FLC* (A) and *VIN3* (B) transcript levels during a vernalization timecourse. RNA was extracted from homozygous plants (Col-FRI, *vin3*, VIN3-GFP WT #3) or from six individual segregating T2 plant lines, grown on media supplemented with the herbicide PPT to select for resistant plants, before vernalization (before cold), at the end of a 6-week cold exposure (6WT0), or 10 days post-cold (6WT10). Data presented are relative to the geometric mean of *UBC* and *PP2A*. *FLC* transcript levels are normalized to *FLC* levels before the cold. Error bars represent standard deviations (*n* = 3 biological replicates).

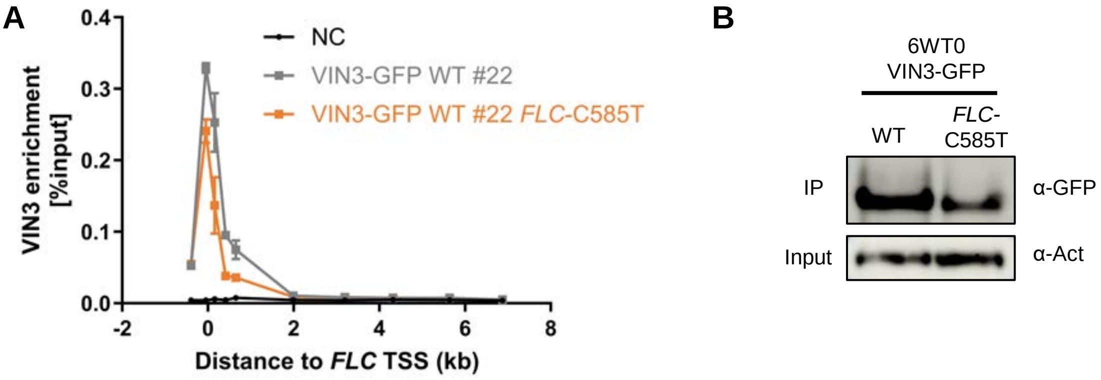


**Fig. S11 Chromatin association of VIN3-GFP in the *FLC-C*585T background.**

(A): ChIP-qPCR showing enrichment of VIN3-GFP across the *FLC* locus (either endogenous or *FLC*-C585T transgene with a point mutation in the first RY motif in intron 1 of *FLC)* in seedlings vernalized for 6 weeks. Note that the small reduction in VIN3-GFP enrichment in the *FLC*-C585T background is attributed to the slightly lower VIN3-GFP levels in this line in comparison to the parental line (Fig. S9B). Non-transgenic Col-FRI plants were used as a negative control sample (NC). Data are shown relative to an input control, error bars represent SEM ($n$ = 2). (B) Immunoblots of α-GFP immunoprecipitates from extracts of vernalized plants bearing the VIN3-GFP transgene in the indicated background. Blots shown are a representative of two replicates.

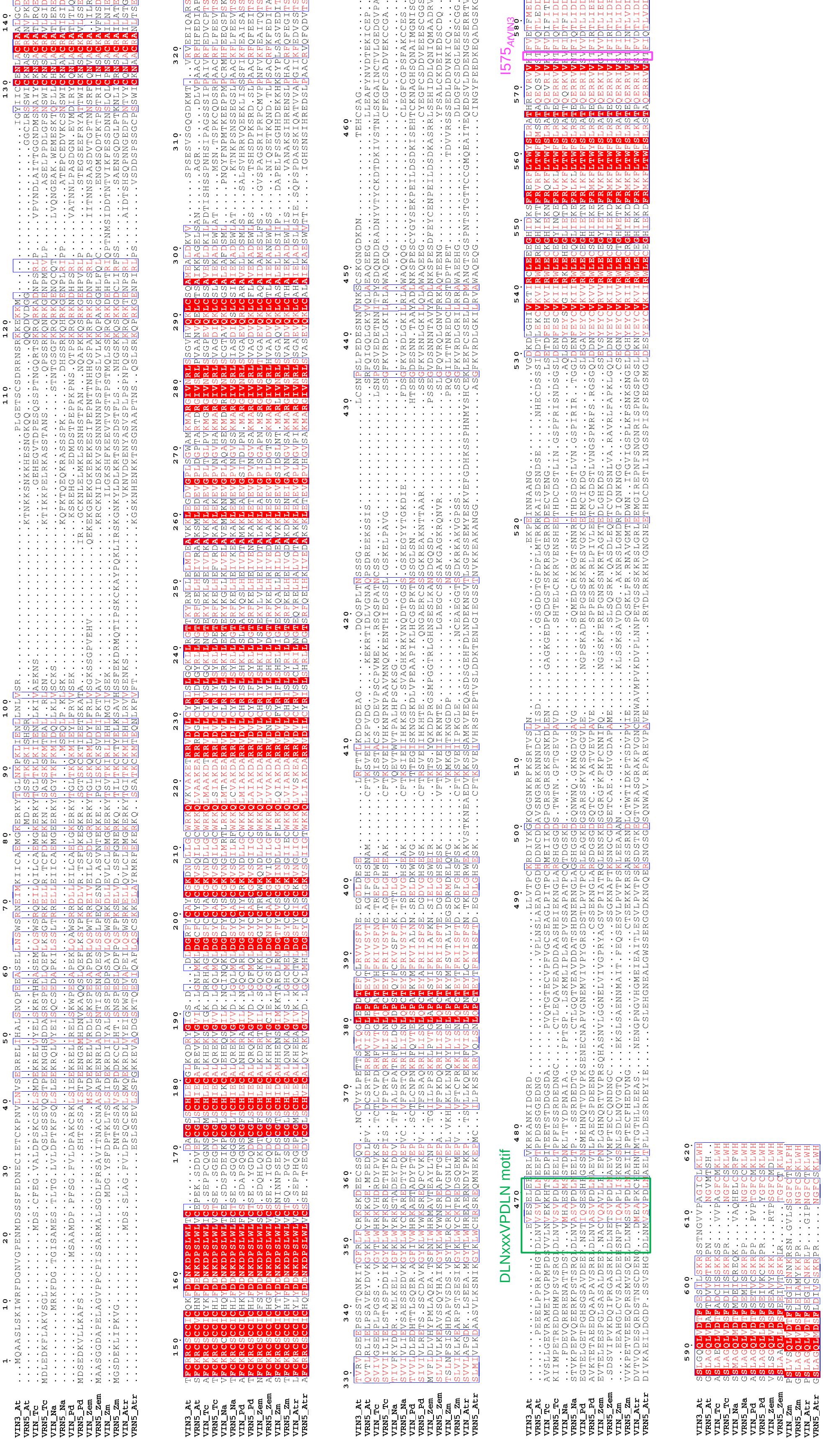


**Fig. S12 Alignment of angiosperm VEL protein sequences (related to Fig. 5B).** VEL protein sequences (VRN5 and VIN3/VEL1 orthologs) from At: *Arabidopsis thaliana*, Tc: *Theobroma cacao*, Na: *Nicotiana attenuata*, Pd: *Phoenix dactylifera*, Zem: *Zea mays*, Zm: *Zostera marina*, Atr: *Amborella trichopoda;* see Supplemental Table S1 for gene IDs). White in red boxes, invariant residues; red in blue frames, similar residues across species; green borders: DLNxxxVPDLN motif that defines VRN5 orthologs[14]; magenta borders: residue differing between VIN3/VEL1 orthologs (hydrophobic amino acids) and VRN5 orthologs (T).
